# TDP-43 is required to build and maintain sarcomeres

**DOI:** 10.64898/2026.06.16.732601

**Authors:** Haileigh Gay, Theodore Ewachiw, Shloka Dhar, Michael Stowell, Bradley Olwin

## Abstract

Skeletal muscle contractile units or sarcomeres require constant maintenance as they are subjected to continuous chemical and mechanical stress. When sarcomere maintenance is disrupted, as occurs in progressive neuromuscular diseases, muscle progressively atrophies, reducing muscle strength and motor control. Transient cytoplasmic ribonucleoprotein aggregates comprised of TDP-43 bound to mRNAs encoding sarcomeric structural proteins (myo-granules) are implicated in building muscle. Ablating TDP-43 in differentiated skeletal muscle causes phenotypes remarkably similar to those of progressive neuromuscular diseases, including muscle atrophy, loss of muscle mass, and aberrantly organized sarcomeres. When injured, differentiated muscle lacking TDP-43 is incapable of repair, failing to build sarcomeres, severely disrupting muscle morphology with fibrotic tissue replacing muscle tissue. TDP-43 is thus required to build and maintain sarcomeres, likely protecting and transporting mRNAs encoding sarcomeric structural proteins in myo-granules.

## Introduction

Skeletal muscle requires continued maintenance and repair to rebuild and replace degraded and damaged contractile proteins that assemble as sarcomeres.^1^ Sarcomeres are comprised of the largest proteins in the human genome, Titin and Nebulin, anchored to alpha-actinin-containing Z-disks, uniformly spaced to maximize muscle force.^2–4^ Although sarcomeres are maintained by frequently replacing protein components, the mechanisms involved are understudied.^5,6^ Moreover, the mechanisms involved in translating Titin and Nebulin, delivering the proteins to the sarcomere, and assembling sarcomeres are not well understood.

Elucidating the mechanisms involved in building and maintaining sarcomeres will likely aid in a more thorough understanding of muscle loss that occurs during aging and in progressive neuromuscular diseases where sarcomeres lose structural alignment and consistent spacing.^7,8^ Whether disrupted sarcomeric structures are responsible for degrading muscle mass, muscle strength, and increasing fibrotic tissue infiltrates or a consequence of neuromuscular disease is not known.^9–11^ Commonly associated with neuromuscular disease are prevalent cytoplasmic mRNA-protein aggregates possessing RNA-binding proteins and sequestered mRNA.^12–15^ The aggregates continuously sequester proteins, propagate to form fibrils, are considered irreversible and theorized to be disease causative.^13,15–17^ However, similarly comprised aggregates or myo-granules, transiently form in the cytoplasm of normal regenerating myofibers, dissipating once sarcomeres are built.^18^ Myo-granules, which consist of mRNAs encoding sarcomeric proteins, the RNA-binding protein TDP-43 and Valosin-containing protein, VCP, transiently appear in the myofiber cytoplasm following a muscle injury and dissipate by ∼10 days post muscle and thus, do not seed irreversible aggregates.^18^

We specifically ablated TDP-43 in differentiated muscle using a mature skeletal muscle-specific promoter to drive Cre recombinase.^19^ Muscles that lack TDP-43 progressively lose strength and atrophy between 2 mo and 6 mo after deleting TDP-43. Sarcomeric structure is disrupted as electron microscopy imaging confirmed sarcomeres lacked consistent alignment and Z-disk length was variable. Upon an induced muscle injury, TDP-43 null muscle was incapable of regenerating, failed to construct sarcomeres, and eventually, significant areas of muscle tissue became fibrotic. We conclude that TDP-43 is essential to maintain sarcomeres and to rebuild sarcomeres following a muscle injury.

## Results

### Muscle-specific inducible Knock-Out of TDP-43

Myo-granules, cytoplasmic aggregates containing TDP-43, transiently appear in the myofiber cytoplasm as skeletal muscle is built, between 3d and 10d following an induced muscle injury.^18^ Transcripts encoding sarcomeric structural proteins are enriched in cytoplasmic myo-granules, suggesting myo-granules are involved in building muscle following an induced muscle injury.^18^ To specifically assess whether TDP-43 is necessary to build muscle, we bred *HSA*^*CreERT2*^ mice with *Tardbp*^*loxp/loxp*^ (TDP-43iKO) to produce an inducible knock out of TDP-43 in differentiated muscle cells committed to terminally differentiate and myonuclei present in myofibers (Fig. 1A). TDP-43 was effectively ablated from myonuclei as 30d post-recombination (Fig. 1B). TDP-43+ myonuclei are readily identified in wild type control mice (*HSA*^*CreERT2*^; *Tardbp*^*+/+*^) mice but virtually undetectable in TDP-43iKO myonuclei (Fig. 1C). When quantified, the percent of TDP-43+ myonuclei was reduced 11-fold from wild type myofibers (Fig. 1D).

**Figure 1:**
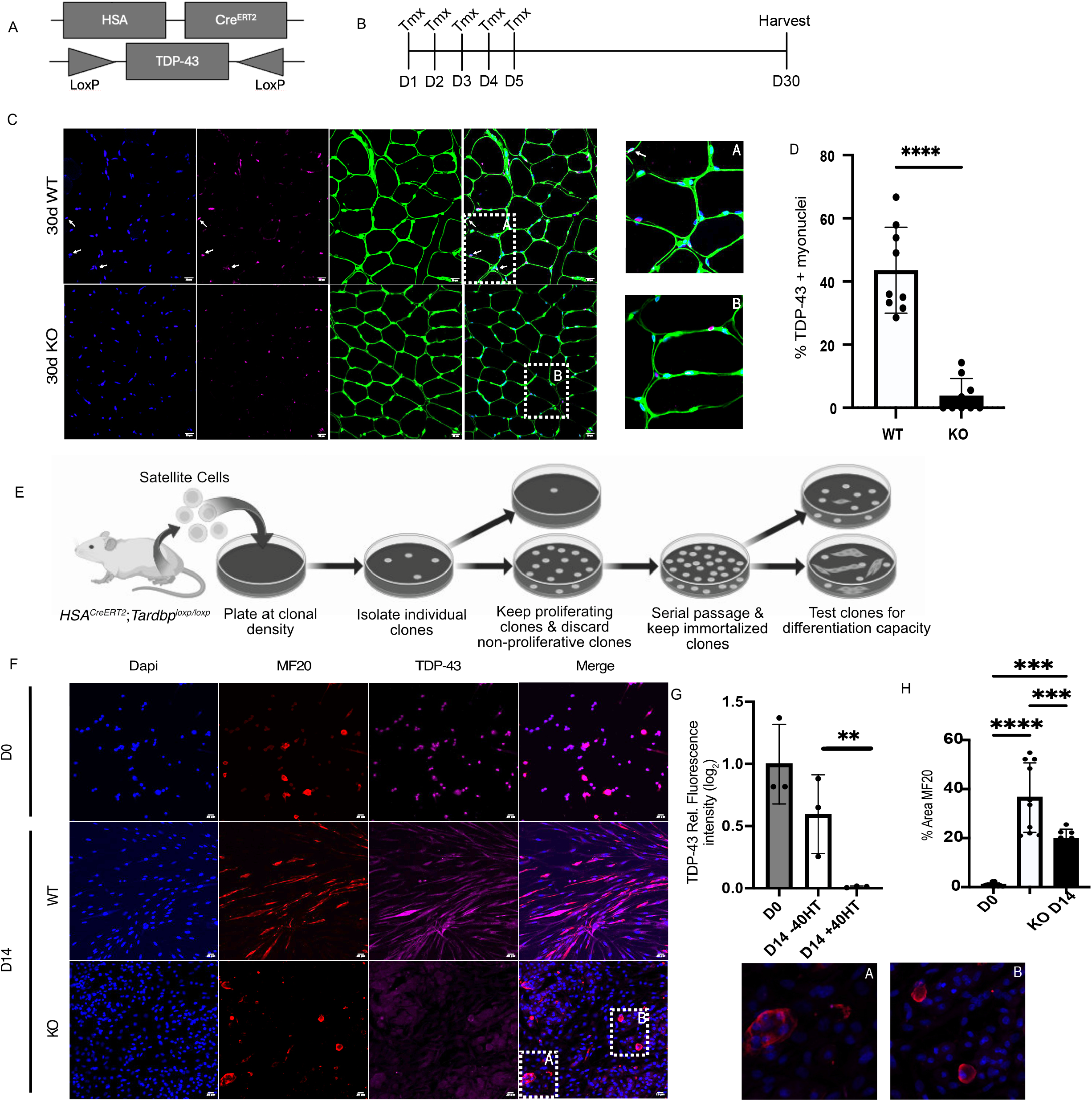
Muscle-specific inducible knock out of TDP-43. **A**. Schematic of TDP-43iKO mouse genetics. **B**. Treatment graphic depicting daily tamoxifen injections over a 5-day period. **C**. Immunoreactivity for TDP-43 in TA muscle sections images of cross-sectioned TA muscle isolated from wild type control mice and TDP-43iKO mice 30d post tamoxifen injection. and TDP-43iKO mice after tamoxifen injections. **D**. TA muscle sections quantified for the fraction of myonuclei that are TDP-43 immunoreactive. n=3 and compared using a two-tailed unpaired t-test. **E**. Schematic of immortalizing cell lines from isolated muscle stem cells. **F**. Immunoreactivity for TDP-43 and MF20 in TDP-43iKO cell line treated with 4-O-hydroxytamoxifen or vehicle. **G**. Quantified relative TDP-43 immunoreactive fluorescent intensities. **H**. Quantified relative MF20 immunoreactive fluorescent intensities. H. MF20+ % area. n=3 and compared via one-way ANOVA

Myonuclei in TA muscle sections likely under-represent the myonuclear population as scoring myonuclei in muscle cross sections fails to reliably quantify all myonuclei. Therefore, to better assess and quantify the efficiency of TDP-43 knockout in differentiated nuclei, we generated an immortalized cell line from isolated stem cells from TDP-43iKO mice (Fig. 1E). Immortalized cells were treated with 4-O-hydroxytamoxifen or a vehicle control, induced to differentiate by removing FGF-2, and reducing serum to 5%. At 14d post-differentiation, the cultures were fixed and assessed for TDP-43 immunoreactivity in muscle-specific myosin (MF-20+) immunoreactive cells (Fig. 1F). When quantified, TDP-43 immunoreactivity was virtually undetectable, and the number of differentiated cells dramatically reduced in TDP-43iKO cells compared to wild type controls (Fig. 1G) where differentiated TDP-43iKO cells appear dysmorphic with spherical multinucleated cells as opposed to typical elongated multinucleated myotubes (Fig. 1F). Thus, TDP-43 is effectively ablated in myonuclei in TDP-43iKO mice and in terminally differentiating cultured cells.

### TDP-43 is required for muscle repair

Cultured cells induced to differentiate fail to form linear, multinucleated myotubes and thus, we asked if injured muscle could repair without TDP-43. TDP-43iKO and wild type mice were injected with tamoxifen prior to BaCl_2_ injury, then harvested at 14 days post-injury (dpi) and 30 dpi (Fig. 2A). TA muscles collected at 30dpi from TDP-43iKO mice were embryonic myosin heavy chain (eMHC) immunoreactive (Fig. 2B) indicating persistent ongoing sarcomerogensis as eMHC is typically replaced by mature muscle isoforms by 10dpi in wild type mice (Fig. 2C). When quantified TDP-43iKO TA muscles possess 35-fold more eMHC immunoreactivity compared to wildtype TA muscle at 30 dpi (Fig. 2E) even though Pax7^+^ cell numbers are indistinguishable between wild type and TDP-43iKO mice (Fig. 2D). Ablating TDP-43 in differentiated muscle does not alter muscle stem cell abundance and thus, no significant differences in numbers of centrally located myonuclei were observed in TDP-43iKO mice compared to wild type mice (Fig. 2E).

**Figure 2:**
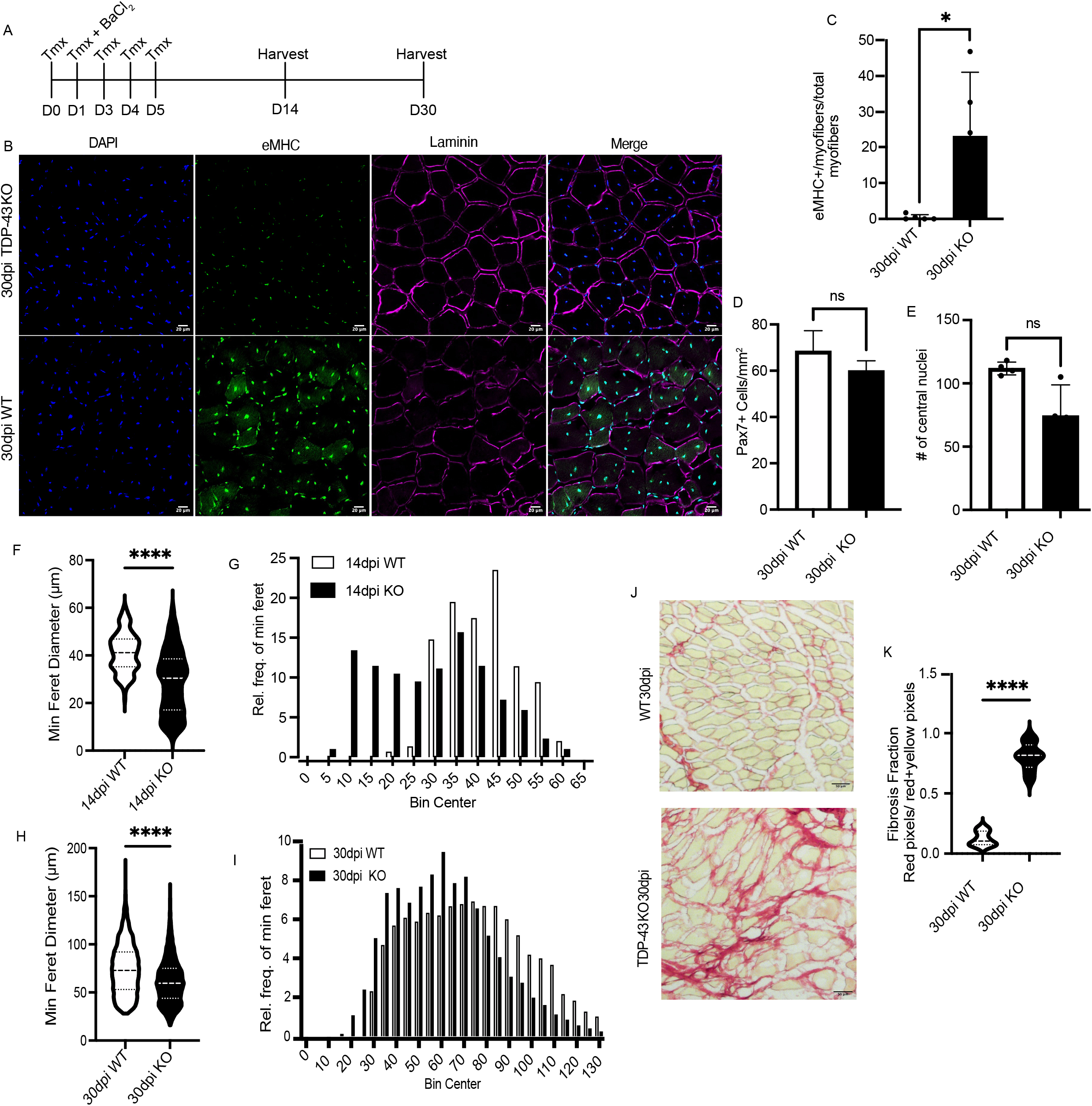
TDP-43 is required for muscle repair. **A**. Treatment graphic showing daily tamoxifen injections beginning before BaCl_2_ injury, day of injury, and 3 days after injury for 14dpi and 30dpi. **B**. Images of 30dpi cross sectioned TA muscles visualized for immunoreactivity of embryonic myosin heavy chain (eMHC) and Laminin. **C**. Pax7+ cells/mm^2^ quantified from TDP-43iKO and wild type mice. **D**. Number of central nuclei quantified in wild type and TDP-43 KO muscle in the presence or absence of TDP-43. **E**. The percentage of myofibers quantified for immunoreactivity to embryonic myosin heavy chain. **F-I**. Minimum feret diameter of 30dpi and 14dpi TA muscle cross sections, respectively. **G**. Picrosirius red histochemical stain of 30dpi cross-sectioned TA muscle in wild type mice and TDP-43iKO mice. **H**. Fibrotic tissue quantified with and without TDP-43 KO using the PSR_quant fiji macro with and without TDP-43. n=3 and comparison done using unpaired t-test.

Muscle morphology was evaluated to determine why eMHC is present at 30dpi in TDP-43iKO TA muscles. TA muscles isolated from TDP-43iKO and wild type mice at 14dpi and 30dpi were assessed for muscle morphometrics and fibrotic tissue. At 14dpi, myofibers from TDP-43iKO mice are significantly smaller than myofibers from wild type mice (Fig. 2F), and the minimum feret diameters are bimodally distributed, with one minimum feret diameter distribution centered at 10-25 mm and a second distribution centered at 25-45 mm (Fig. 2G). At 30dpi, myofibers from TDP-43iKO mice were significantly smaller than in wild type mice (Fig. 2H, I). Myofiber minimum feret diameters from TDP-43iKO mice vary more widely in diameter than myofibers from wild type mice at 14 and 30dpi (Supplemental Fig. 2A, B). The subset of larger minimum feret diameter myofibers likely represents myofibers that were either not injured or received a minor injury and thus did not regenerate, maintaining their original minimum feret diameter. The smaller minimum feret myofiber diameter at 10-25mm likely arises from the failure of muscle to regenerate following the induced injury. Collagen I and III, identified by picrosirius red, infiltrate muscle extracellular space in TDP-43iKO mouse muscle where myofibers fail to regenerate (Fig. 2J). Isolated TA muscle from TDP-43iKO contains 6.5-fold more picrosirius red area compared to wild type TA mouse (Fig. 2K). Nearly 80% of the TA muscle area is picrosirius red in TDP-43iKO mice, whereas in wild type mice only 10-20% of the TA muscle is picrosirius red+ identifying blood vessels, tendons, and muscle spindle cells (Fig. 2J, K). The 60% increase of picrosirius red staining in TDP-43iKO mouse muscle likely represents fibrotic tissue infiltrating muscle, where myofibers fail to regenerate to provide structural support.

### Cellular architecture and morphology degrade without TDP-43 following injury

TDP-43iKO mice fail to regenerate following an induced muscle injury, replacing myofibers with fibrotic tissue. To better analyze whether loss of TDP-43 affected sarcomeres, we visualized sarcomeres using transmission electron microscopy (TEM) of extensor digitorum longus (EDL) muscle longitudinal sections at 30dpi from TDP-43iKO mice. The protein content of sarcomeres is visualized by heavy metal staining on electron micrographs (Fig 3A). The Z-disks in TDP-43iKO EDL muscle are severely misaligned when compared to wild type EDL muscle, suggesting sarcomeres are dysfunctional (Fig. 3A). Mitochondria and vesicle infiltrates appear in between Z-disks of TDP-43iKO EDL muscles (Fig. 3A) and Z-disk average length differs significantly from wild type muscle (Fig. 3B). Z-disks anchor thin and thick filaments during repair and as Z-disks are hindered from forming the sarcomere is likely perturbed and nonfunctional. Thus, TDP-43 is essential for muscle to build functional sarcomeres capable of adequate contractile force.

**Figure 3:**
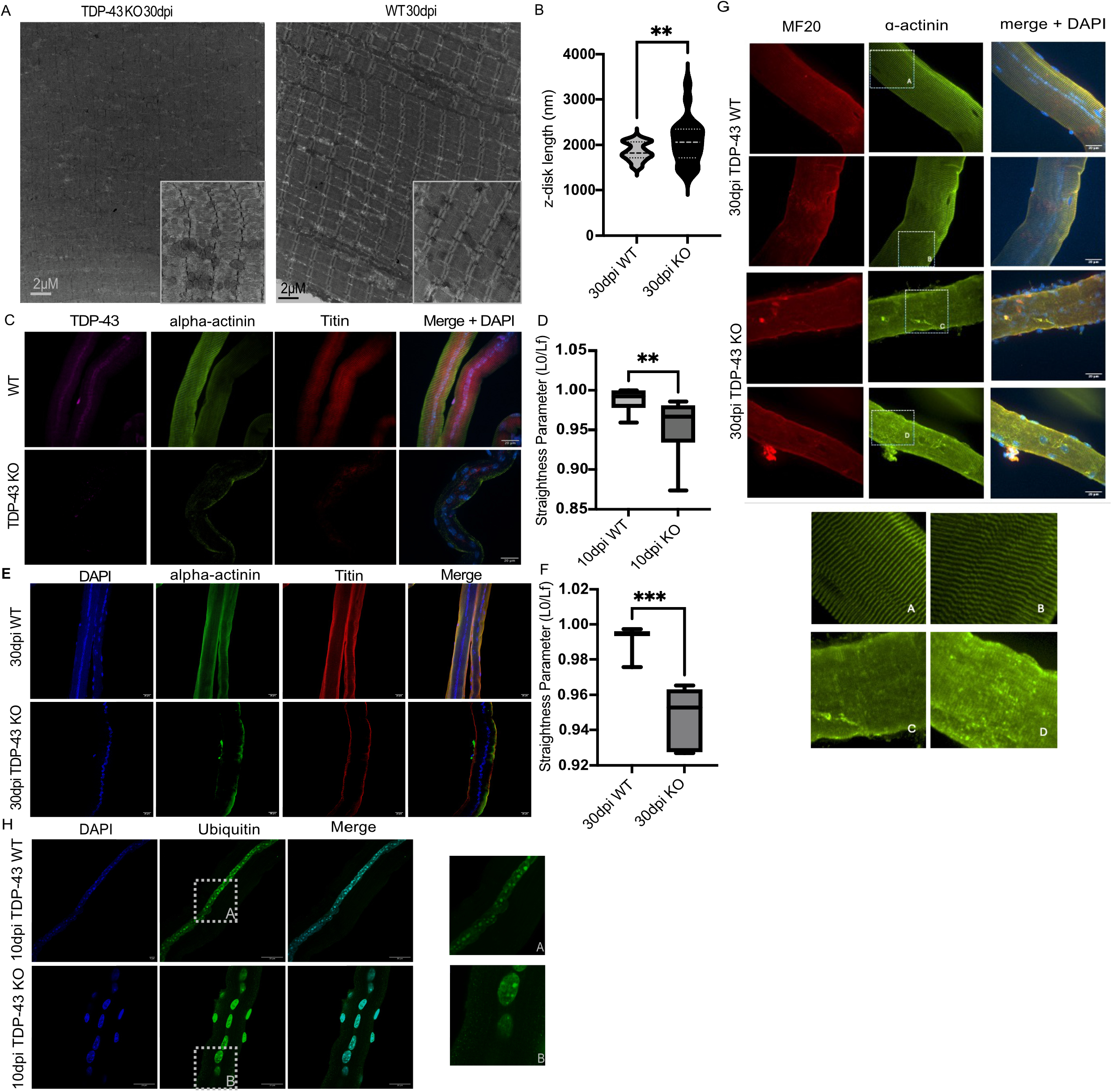
TDP-43 is necessary to build sarcomeres. **A**. Transmission electron tomographs of longitudinally sectioned EDL muscles 30dpi with and without TDP-43 KO. **B**. Z-disk alignments quantified in 30dpi muscle in TDP-43iKO and in wild type mice are significantly different. **C**. Myofibers isolated from 10dpi TDP-43iKO and wild type mice treated with tamoxifen and assessed for alpha-actinin and Titin immunoreactivity. **D**. Myofiber linearity quantified at 10dpi quantified using NeuronJ Fiji plugin. **E**. Myofibers isolated from 30dpi TDP-43iKO and wild type mice treated with tamoxifen and assessed for alpha-actinin and Titin immunoreactivity. **F**. Myofibers linearity quantified at 30dpi. n=3 comparison done using a two-tailed unpaired t-test. **G**. Myofibers isolated from TDP-43iKO and wild type mice at 30dpi and assessed for MF20 and alpha-actinin immunoreactivity (magnified images from areas highlighted with a dotted line box). **H**. Myofibers isolated from TDP-43iKO and wild type mice at 30dpi and assayed for immunoreactive ubiquitin protein (magnified images from areas highlighted with a dotted line box).

Myofibers isolated from TDP-43iKO mice at 10dpi display aberrant morphology compared to wild type myofibers; in the absence of TDP-43 myofibers do not maintain a consistent shape when isolated and collapse (Fig. 3C). The earliest time point to consistently isolate myofibers following an injury is 10dpi. Measurements of plasma membrane linearity of myofibers are significantly reduced in TDP-43iKO mice when compared to myofibers from wild type mice indicating a lack of structural integrity. Wild type myofiber plasma membrane linearity varies from 0.95-1 and TDP-43iKO myofiber plasma membrane linearity varies from 0.87-0.96 (Fig. 3D). At 30dpi, the linearity of myofibers from TDP-43iKO mice does not improve and diverges more significantly from wild type myofiber linearity than at 10dpi (Fig. 3F). Myofibers isolated from TDP-43iKO mice 30dpi contained proteinaceous cytoplasmic puncta, immunoreactive for alpha-actinin that were absent in myofibers from wild type mice with fully formed and striated sarcomeres (Fig 3G). Protein-mRNA aggregates are present during muscle regeneration following BaCl_2_-mediated injury in wild type mice, peaking 5dpi, and undetectable at 10dpi.^20^ Myofibers isolated at 10dpi from TDP-43iKO mice contained cytoplasmic aggregates immunoreactive for ubiquitin protein, a canonical aggregate marker (Fig. 3H). Ubiquitinated aggregates localized at the edges of TDP-43iKO myofibers, where minimal immunoreactivity for Titin and alpha-actinin protein was localized; aggregates are undetectable in myofibers from wild type mice.

Myofibers incapable of repair, likely do not survive the myofiber isolation and are underrepresented in myofiber imaging and quantification. Surviving myofibers isolated from TDP-43iKO mice do not contain detectable sarcomere structures and cellular morphology is aberrant compared to wild type myofibers. Myofibers without TDP-43 are thus incapable of building sarcomeres, causing myofibers to collapse and disrupting linear cell morphology, thereby maintaining persistent aggregates.

### Muscle progressively atrophies and loses strength without TDP-43

Wild type and TDP-43iKO mice were injected with tamoxifen for 5 consecutive days to ablate TDP-43 and for 3 consecutive days in each following month before harvesting TA muscles at 6 months of TDP-43 loss (Fig. 4A). When TDP-43 was ablated, myofiber minimum feret diameter decreased by 1.4-fold when compared to wild type TA muscles (Fig. 4B, C). Despite a 10-fold increase in centrally located myonuclei in TDP-43iKO TA muscles compared to wild type TA muscles (Fig. 4D), the TA muscle weight decreased by 1.5-fold when TDP-43 was ablated for 6 months (Fig. 4E). Muscle mass was reduced and myofiber diameter declined in TDP-43iKO TA muscles despite the extensive increase in centrally located myonuclei reflecting satellite cell fusion. Ablating TDP-43 causes a progressive loss of muscle tissue, reflected in a loss of body weight over the 6-month period (Fig. 4F). Even though satellite cells extensively fused, TDP-43iKO mice undergo severe muscle atrophy resulting in a dramatic loss of body weight.

**Figure 4:**
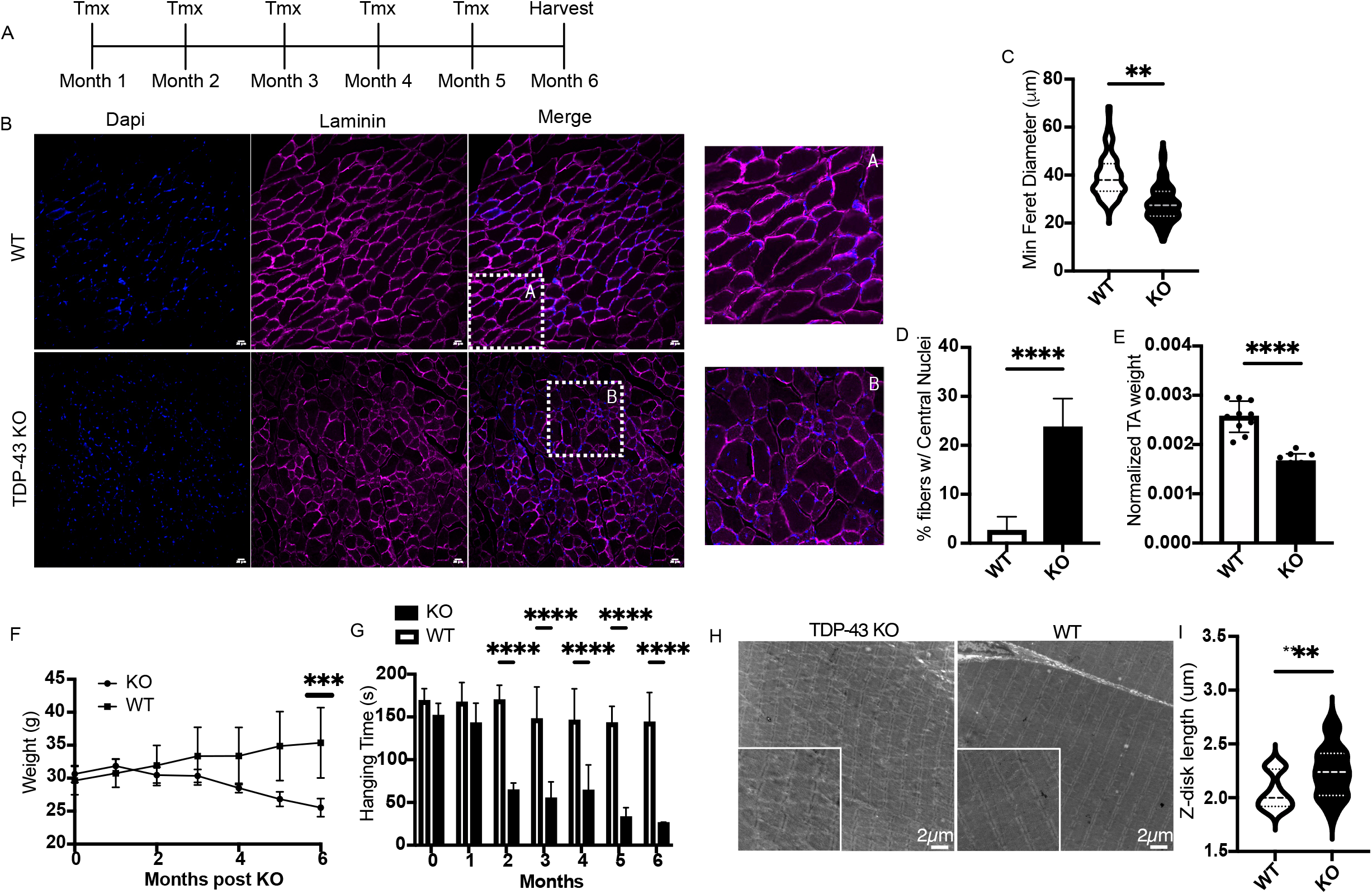
TDP-43 is required for muscle strength and structure. **A**. Treatment graphic depicting daily tamoxifen injections and monthly 3-day injections for 6 months. **B**. TA muscle cross sections from wild type and TDP-43iKO mice treated as shown in A and assessed for laminin immunoreactivity. **C**. Myofiber minimum feret diameters quantified from TA muscles in wild type and TDP-43iKO mice. **D**. The percent myofibers with centrally located myonuclei quantified in TDP-43iKO mice and wild type mice. **E**. TA muscle weights quantified for TDP-43iKO mice and wild type mice. **F**. Hanging wire test for TDP-43iKO mice and wild type mice. **G**. Total mouse body weight by month for TDP-43iKO mice and wild type mice. N=3 and compared using a 2-way ANOVA. **H**. Electron microscopy images of cross-sectioned EDL muscle in TDP-43iKO mice and wild type mice. **I**. Z-disk length quantified in muscle from TDP-43iKO mice and wild type mice following 6 mo of tamoxifen. All tests n=3 and compared using a two-tailed unpaired t-test unless stated otherwise.

A loss of muscle mass in TDP-43iKO mice should reflect compromised muscle function in TDP-43iKO mice compared to wild type mice. TDP-43iKO and wild type mouse muscle strength was assessed by a longitudinal hanging wire test. TDP-43iKO mice rapidly lost strength between 1 mo and 2 mo of TDP-43 loss, maintaining one-half of their hanging time until 5 mo post ablating TDP-43 where another 2-fold loss in hanging wire time occurred (Fig. 4G). EDL muscles from TDP-43iKO and wild type mice 6 mo after ablating TDP-43 were imaged with electron tomography. Z-disks in TDP-43iKO mice appear aberrant with misalignment of Z lines and a lack of linearity compared to wild type EDL muscle (Fig 4H). The Z-disk length in TDP-43iKO mice is longer and more variable than those in wild type mice (Fig. 4I). In addition to muscle atrophy, the remaining sarcomeres appear aberrant when TDP-43 is ablated for 6 mo, indicating a failure to maintain as well as regenerate muscle.

## Discussion

Ablating TDP-43 from differentiated skeletal muscle myonuclei caused a rapid and progressive loss of skeletal muscle mass, accompanied by disorganized sarcomeres and a striking loss of strength. Therefore, TDP-43 is essential to maintain skeletal muscle integrity. Since skeletal muscle is continuously contracting, causing mechanical and chemical stress, sarcomere proteins are constantly replaced.^21^ Myo-granules, transient cytoplasmic aggregates, contain TDP-43, which binds mRNAs encoding sarcomeric structural proteins.^18^ If TDP-43 organizes and protects sarcomeric structural mRNAs^18^ then loss of TDP-43 may prevent sarcomere maintenance, consistent with the observed loss in muscle mass when TDP-43 was ablated.

To directly ask if skeletal muscle can be built in the absence of TDP-43, we recombined TDP-43iKO mice and then induced a muscle injury in the TA muscle. Within the first 4 days following an injury, muscle stem cells expand and fuse to form small myofibers with centrally located nuclei that then grow by additional cells fusing to produce peripheral myonuclei. The first myosin to appear, eMHC, is detected soon after centrally nucleated myofibers form at 5dpi and is lost by 10 dpi when eMHC is replaced by more mature myosin isoforms. Myo-granule appearance and disappearance in the myofiber cytoplasm coincides with detectable eMHC protein.^18^ In TDP-43iKO mice, eMHC persists to at least 30dpi, suggesting continuous attempted muscle building well beyond the wild type time frame of 10 dpi. Building muscle appears inefficient in TDP-43iKO since myofibers from injured TA muscles were eMHC immunoreactive, significantly smaller, and varied more broadly in minimum feret diameter than myofibers from wild type mouse TA muscles at 30 dpi. Accompanying the smaller regenerated TA muscles in TDP-43iKO mice was extensive fibrotic tissue, further indicative of a failure to generate muscle tissue. The observed failure to regenerate muscle was not due to deficits in the satellite cells as satellite cell numbers were indistinguishable when comparing TA muscles from TDP-43iKO and wild type mice at 30 dpi. Nor was the failure to regenerate due to an inability of satellite cells to fuse, as the numbers of central myonuclei in injured TA muscle of TDP-43iKO mice and wild type mice were indistinguishable. Thus, we believe that the failure to regenerate muscle in TDP-43iKO mice following an injury arises from the loss of TDP-43 in differentiated muscle and likely occurs because sarcomeres cannot be built.

Sarcomeres align in uniform units with Titin protein serving as a molecular ruler to precisely distance and align adjacent Z-disks anchoring thick and thin filaments to maximize muscle contraction.^4,22,23^ TDP-43iKO mice appear incapable of building sarcomeres as Z-disks broadly vary in distance from one another and are not aligned with adjacent Z-disks, likely preventing muscle from contracting. Lipid micelles and mitochondria infiltrate sarcomeres, likely preventing Z-disks from contracting. TDP-43iKO mice fail to generate striated sarcomeres containing titin and alpha-actinin, strongly implicating TDP-43 in building sarcomeres. The lack of myofibrillar structure affects myofiber cellular morphology in TDP-43iKO mice, as isolated myofibers are aberrant, exhibiting kinks, bends and misaligned and non-uniformly dispersed myonuclei. TDP-43iKO mice are delayed in building sarcomeres as myofibers at 30 dpi do not form regularly spaced sarcomeres and have minimal immunoreactive alpha-actinin and titin at the edges of the myofiber, suggesting that the absence of sarcomeres arises from the failure to synthesize sarcomere proteins and organize sarcomeres. The persistent eMHC immunoreactivity and extensive fibrosis in TDP-43 null muscle are consistent with the failure to build sarcomeres. Thus, we conclude that TDP-43 is essential to build muscle and the loss of TDP-43 affects myo-granule function, likely affecting sarcomeric structural mRNA half-lives and mislocalizing myo-granules, resulting in a failure of sarcomerogenesis.

Cytoplasmic TDP-43 aggregates are prominent in the muscle of neuromuscular disease patients, where progressive loss of skeletal muscle consistently results in severe muscle atrophy and loss of strength.^12,24,25^ Muscle cannot be rebuilt following an injury in TDP-43iKO mice, and thus, we asked if TDP-43 is required to maintain skeletal muscle by ablating TDP-43 for 6 months in differentiated muscle. The striking progressive losses of muscle strength, mouse weight, and muscle mass in recombined TDP-43iKO mice demonstrate a requirement for TDP-43 to maintain muscle tissue. Accompanying the loss of muscle strength are sarcomere phenotypes like those that occur following an induced muscle injury. Surprisingly, at 6 mo post-ablating TDP-43, central myonuclei increase 20-fold, reflecting a major increase in satellite cell fusion. The loss of TDP-43 is most likely affecting myo-granule function, negatively impacting the half-lives and subcellular localization of mRNAs encoding sarcomeric structural proteins, and exacerbating cell stress as sarcomeres cannot be maintained. Since TDP-43iKO mice lost significant body weight, likely due to muscle atrophy arising from TDP-43 loss, TDP-43 appears essential for myo-granules to maintain and repair sarcomeres supporting muscle homeostasis during normal muscle function.

Skeletal muscle phenotypes for progressive neuromuscular diseases are strikingly similar to skeletal muscle phenotypes in mouse muscle lacking TDP-43, including persistent protein aggregates in the cytoplasm, failure to maintain and rebuild sarcomeres, major increases in satellite cell fusion resulting in the deposition of central myonuclei, loss of muscle function, and muscle atrophy. Persistent cytoplasmic aggregates present in TDP-43iKO mouse muscle and in mouse models for neuromuscular diseases, as well as neuromuscular disease patients, may be a direct consequence of attempting to rebuild muscle where the genetic deficits affect myo-granule function, preventing maintenance and replacement of sarcomeres. Thus, simply dissolving the persistent aggregates will not improve disease phenotypes, as the underlying function of myo-granules in wild type muscle tissue has been compromised. Attempts to rebuild muscle by engaging satellite cells and satellite cell fusion further aggravate the phenotypes as the attempted regenerative response initiates transient myo-granule assembly that may promote irreversible aggregate formation.^20^ Thus, failure of myo-granules to function would directly affect the ability of muscle to form and maintain sarcomeres rather than the aggregates non-specifically interfering with muscle tissue.

The requirement for TDP-43 in maintaining and repairing muscle recapitulates aging and disease-related muscle atrophy, loss of strength and endurance, and increased fibrosis. Irreversible aggregates in disease may sequester TDP-43 and prevent myo-granule-mediated RNA processing and trafficking necessary to rebuild and maintain muscle fibers and sarcomeres. When TDP-43 is sequestered or prevented from functioning, aggregates persist and propagate, causing disease onset and progression. The mechanism by which TDP-43 binds, packages, and transports mRNA for myo-granule function remains elusive; thus, better understanding physiological maintenance and repair mechanisms may aid in understanding how aggregates irreversibly form in neuromuscular disorders.

## METHODS

### Mice

Mice were bred and housed according to National Institute of Health (NIH) guidelines for the ethical treatment of animals in a pathogen-free facility at the University of Colorado at Boulder. The University of Colorado Institutional Animal Care and Use Committee (IACUC) approved all animal protocols and procedures to ensure all studies complied with all ethical regulations. Mice used were C57BL6 (Jackson Labs Stock No. 000664), *HSA*^*CreERT2*^ (039097), *Tardbp*^*loxp/loxp*^ (017589), mice. Crossing *Tardbp*^*loxp/loxp*^ mice into *HSA*^*CreERT2*^ generated conditional *Tardbp*^*loxp/+*^ or *Tardbp*^*loxp/loxp*^ mice. Crossing conditional *Tardbp*^*loxp/+*^ or *Tardbp*^*loxp/loxp*^ mice with *HSA*^*CreERT2*^ mice generated *HSA*^*CreERT2*^;*Tardbp*^*loxp/+*^ mice (TDP-43iKO). Tibialis anterior muscles were isolated from mice between. Control mice were randomly assigned and were sex and age matched to the mice and crosses outlined above. Sample sizes were set at *n* = 3. Each mouse used in this study was genotyped by Transnetyx (Cordova, TN).

### Mouse injuries and tamoxifen delivery

Mice were anesthetized with isoflurane followed by injection with 50µL of 1.2% BaCl2 in sterile saline into the TA muscle, which injures the extensor digitorious longus along with the TA muscles. Tamoxifen (Sigma-Aldrich) was resuspended in corn oil (Sigma-Aldrich) and administered by intraperitoneal injection at a volume of 0.075 mg of tamoxifen per gram of mouse weight.

### Mouse Hanging Wire Test

Mice were given three attempts to grip onto a constructed twelve-gauge circular rolling wire with all four paws for a maximum of 180 seconds. The time before falling for each attempt was recorded. Sample sizes were set at *n* =3. Data was analyzed through a two-way Anova.

### Cell harvest and cell line establishment

Gastrocnemius, extensor digitorum longus, tibialis anterior and all other lower hindlimb muscles were dissected from wild type mice. The muscle groups from both hindlimbs were separated and digested in 20 ml of Ham’s F12 media (Gibco) supplemented with 0.8mM CaCl_2_ (Fisher), penicillin and streptomycin (Gibco) and 400U/mL collagenase (Worthington) for 1hr at 37 °C vortexing for 30 sec every 10 min. Collagenase digestion was quenched immediately and cells were passed through three strainers of 100μm, 70μm, and 40μm (BD Falcon). The flow through was centrifuged at 1500×g for 5 min and the cell pellets were re-suspended in Ham’s F-12C. Cells were counted and plated at a range of low densities in 10 ml growth medium at 37 °C supplemented with 50nM FGF-2. Colonies of myoblasts formed by four days of culture and were expanded by passaging with 0.25% trypsin-EDTA (Sigma-Aldrich).

### Myofiber Isolation and immunostaining

The EDL muscles were dissected and placed into 400 U/mL collagenase at 37°C for 1.5h with shaking and then transferred into Ham’s F-12C and 15% horse serum to inactivate the collagenase. Individual EDL myofibers were separated using a glass pipet and immediately fixed using 4% paraformaldehyde for 10 min at room temperature and stored in PBS at 4°C. Myofibers were then permeabilized with 0.25% Triton-X100 in PBS (phosphate buffered saline without calcium or magnesium pH 7.4), containing 3% bovine serum albumin (Sigma) for 1hr at room temperature. The intact myofibers were incubated with mouse anti-alpha-actinin (Abcam) at 1:400, rabbit anti-titin (Proteintech) at 1:200, and rabbit anti-ubiquitin (Cytoskeleton) at 1:1000 diluted with 0.125% Triton-X100 in PBS at room temperature for 1h followed by three washes in PBS. Alexa secondary antibodies included donkey anti-rabbit 488 and 647, goat anti-mouse IgG2a 647, and goat anti-mouse IgG1 488, 555, and 647 (Thermo) were used at a dilution of 1:1,000 incubated with Dapi (1ug/mL) for 1hr at room temperature. Myofibers were washed 3 times in PBS and then mounted in Mowiol.

### Detection of antigens in tissue sections and cell culture

Tibialis anterior muscles were dissected, fixed on ice for 2 h with 4% paraformaldehyde, and transferred to phosphate-buffered saline (PBS) with 30% sucrose at 4°C overnight. Muscle was mounted in OCT (Tissue-Tek) and cryo-sectioning was performed on a Leica cryostat to generate 10-μm thick sections. Tissues and sections were stored at −80°C until staining. Tissue sections were post-fixed in 4% paraformaldehyde for 10 min at room temperature and washed three times for 5 min in PBS at room temperature. Immunodetection of PAX7 required heat-induced epitope antigen retrieval, for which post-fixed slides were placed in citrate buffer, pH 6.0, and subjected to 6 min of high pressure-cooking in a Cuisinart model CPC-600 pressure cooker. For immunodetection, tissue sections were permeabilized with 0.25% Triton X-100 (Sigma-Aldrich) in PBS containing 2% bovine serum albumin (BSA) (Sigma-Aldrich) for 60 min at room temperature following by incubation with a primary antibody at 4 °C overnight and by incubation with a secondary antibody at room temperature for 1 h. Primary antibodies included mouse anti-PAX7 (Developmental Studies Hybridoma Bank) at 1:750, rabbit anti-TDP-43 (ProteinTech) at 1:200, mouse anti-TDP-43 (Abcam) at 1:200, mouse anti-Panmyosin (DSHB), rabbit anti-titin (Proteintech), rat anti-laminin (Sigma-Aldrich), and mouse anti-alpha actinin (Abcam) at 1:800. Alexa secondary antibodies included donkey anti-rabbit 488 and 647, goat anti-mouse IgG2a 647, and goat anti-mouse IgG1 488, 555, and 647 (Thermo) were used at a dilution of 1:1,000. Sections were incubated with 1 μg ml−1 DAPI for 10 min at room temperature then mounted in Mowiol supplemented with DABCO (Sigma-Aldrich) or ProLong Gold (Thermo) as an anti-fade agent.

### Picrosirius Red Staining

TA muscle sections were prepared as described above and stored at -80°C until staining was performed. Tissue was post-fixed in Bouin’s solution (Sigma-Aldrich) for 1 h at 56°C. Slides were then rinsed under tap water until fixative was removed and Picrosirius Red (American Master Tech) was added for 1 h at room temperature. Slides were then rinsed twice in 0.5% acetic acid (J.T. Baker) for one minute each. Tissue sections were then dehydrated stepwise in 50%, 75%, and 100% ethanol (Sigma-Aldrich) for 30 seconds each. Slides were then rinsed twice in xylene (VWR) for 1 minute and left to dry at room temperature before mounting with permount (Thermo).

### Microscopy and image analysis

Images of immunoreactive tissue and cells were captured on a Nikon A1 laser scanning with resonance and super-resolution where objectives used were 20x/.75NA Plan Apo, 40x/0.95NA Plan Apo, and a 100x/1.49NA OIL SR Apo. For imaging of immunohistochemical stains of picrosirius red stain, a Nikon TiU widefield with RGB color brightfield was used with a 4x 0.13NA Plan Fluor WD objective. All images were processed using Fiji ImageJ. Confocal stacks were projected as maximum intensity projections for each channel and merged to form a single image. Brightness and contrast were adjusted for the image when necessary.

### Transmission electron microscopy

Extensor digitorum longus muscles were dissected as described above and fixed in a 2.5% Glutaraldehyde / 4% PFA fixative in 0.1M NaCaC buffer. The EDL muscles were subsequently sectioned in half and stained using osmium tetroxide, dehydrated through an ethanol series, infiltrated in Epon 812, and polymerized. Sectioned muscle was imaged using a FEI tecnai T12 spririt 120kv LaB6 filament TEM.

## Author Contributions

Conceptualization: HTG, TEE, BBO

Methodology: HTG, TEE, BBO

Investigation: HTG, TEE, SD

Visualization: HTG, TEE, SD

Supervision: BBO, MS

Writing—original draft: HTG, BBO

Writing—review & editing: HTG, BBO

