## Supplementary Figures for "TDP-43 is required to build and maintain sarcomeres"

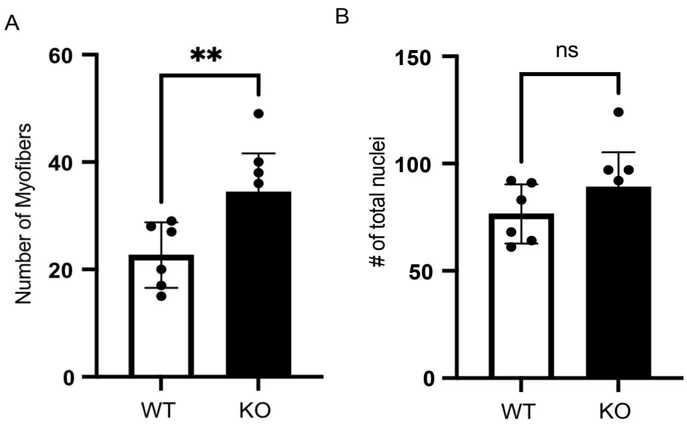
A
Number of Myofibers
60
40
20
0
**
WT
KO
B
### of total nuclei
150
100
50
0
ns
WT
KO

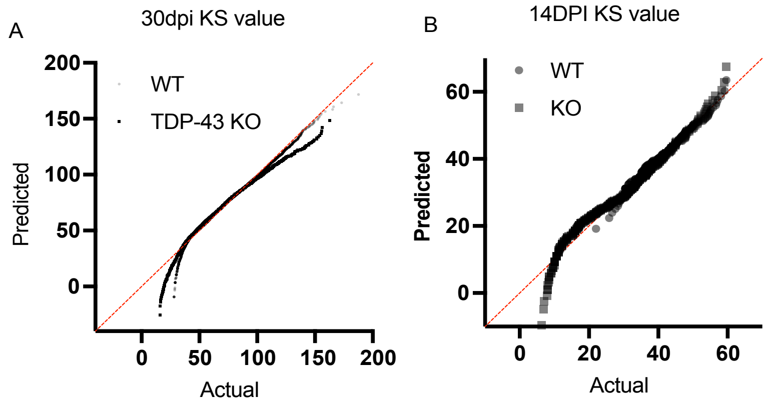
A
30dpi KS value
WT
TDP-43 KO
Predicted
Actual
200
150
100
50
0
0
50
100
150
200
B
14DPI KS value
WT
KO
Predicted
Actual
60
40
20
0
0
20
40
60

#### Supplementary Figures

##### Supplementary Figure 1

A. Myofiber abundance quantified in TDP-43iKO and wild type mice after a 30-day recombination period. B. Number of total nuclei quantified in TDP-43iKO and wild type mouse muscle.

##### Supplementary Figure 2

A-B. Kolmogorov-Smirnov test for 30dpi and 14dpi min feret diameter distributions respectively.
